# Characterisation of pouch secretions from breeding Tasmanian devils

**DOI:** 10.64898/2025.11.30.691433

**Authors:** Manujaya W. Jayamanna Mohottige, Emma Peel, Angéla Juhász, Mitchell G. Nye-Wood, Katherine Belov, Michelle L. Colgrave, Carolyn J. Hogg

**Author notes:** **Corresponding author**: Professor Carolyn Hogg, RMC Gunn Building (B19), The University of Sydney, Sydney, NSW, 2006, Australia.

## Abstract

Tasmanian devils (*Sarcophilus harrisii*), like all marsupials, give birth to altricial young. Here we employ proteomic analysis to identify components of devil pouch secretions (red oil) that may contribute to offspring survival and development. Proteins were extracted from 5 samples and analysed by liquid chromatography-tandem mass spectrometry. Peptide-level evidence revealed proteins involved in a diverse set of immune pathways, including those mediating iron-ion transport, defence responses to bacteria, innate immune responses, and antigen processing. A core set of 200 proteins was identified across at least three samples, 63 of which were associated with antimicrobial or immunoregulatory functions. These included immunoglobulins, components of complement and coagulation cascade, and antimicrobial proteins and peptides. For the first time, these findings highlight the Tasmanian devil’s red oil secretions as a source of immune proteins likely contributing to microbiome restructuring during lactation. Moreover, the data indicate that these proteins may act synergistically in pathways triggered by immune challenges or physiological stress.

**In brief:** Microbial shifts in the Tasmanian devil pouch are critical for establishing a protective environment for the developing joey. This study confirms that Tasmanian devil red oil contains a diverse array of immune proteins that likely contribute to these microbial shifts.

## Introduction

Marsupials are a distinct lineage of mammals’ native to Australia and South America that diverged from eutherian mammals more than 150 million years ago (Luo *et al*. 2011). Their reproductive strategy is characterised by a short gestation, typically ranging from 12 to 38 days, followed by birth of altricial young (Tyndale-Biscoe & Renfree 1987). Postnatal development then continues in the maternal pouch during an extended lactation phase, that functionally parallels the role of the eutherian placenta (Tyndale-Biscoe & Renfree 1987). Unlike eutherians, marsupials are immunologically naïve at birth and lack all mature immune tissues and cells, some of which take up to four months to develop (Borthwick *et al*. 2014). Because the pouch is non-sterile and harbours diverse bacterial communities (Weiss *et al*. 2021, Maidment & Eisenhofer 2023, Ockert *et al*. 2024), marsupials have evolved various protective mechanisms including passive immune transfer via milk and the rapid activation of pouch young innate immunity (Edwards *et al*. 2012).

The Tasmanian devil (*Sarcophilus harrisii*) is the largest extant marsupial carnivore and is endemic to the island state of Tasmania, Australia (**Fig. 1**). Females produce ∼30 embryos per breeding event but possess only four functional teats in their backward facing pouch, resulting in a maximum litter size of four offspring per year (Guiler 1970). The breeding season is from March to June, with joeys attached to the teat for four months before remaining in the den until December-January when they disperse (Guiler 1970, Keeley *et al*. 2017). The Tasmanian devil pouch undergoes striking morphological changes across the reproductive cycle (**Fig. 1D**). Prior to and during mating, it develops a “red oil” and “lipstick ring” appearance (**Fig. 1C**; Hesterman *et al*. 2008). The red oil starts to diminish as the young joeys attach to the teats and start to grow (Hesterman *et al*. 2008).

**Figure 1.**
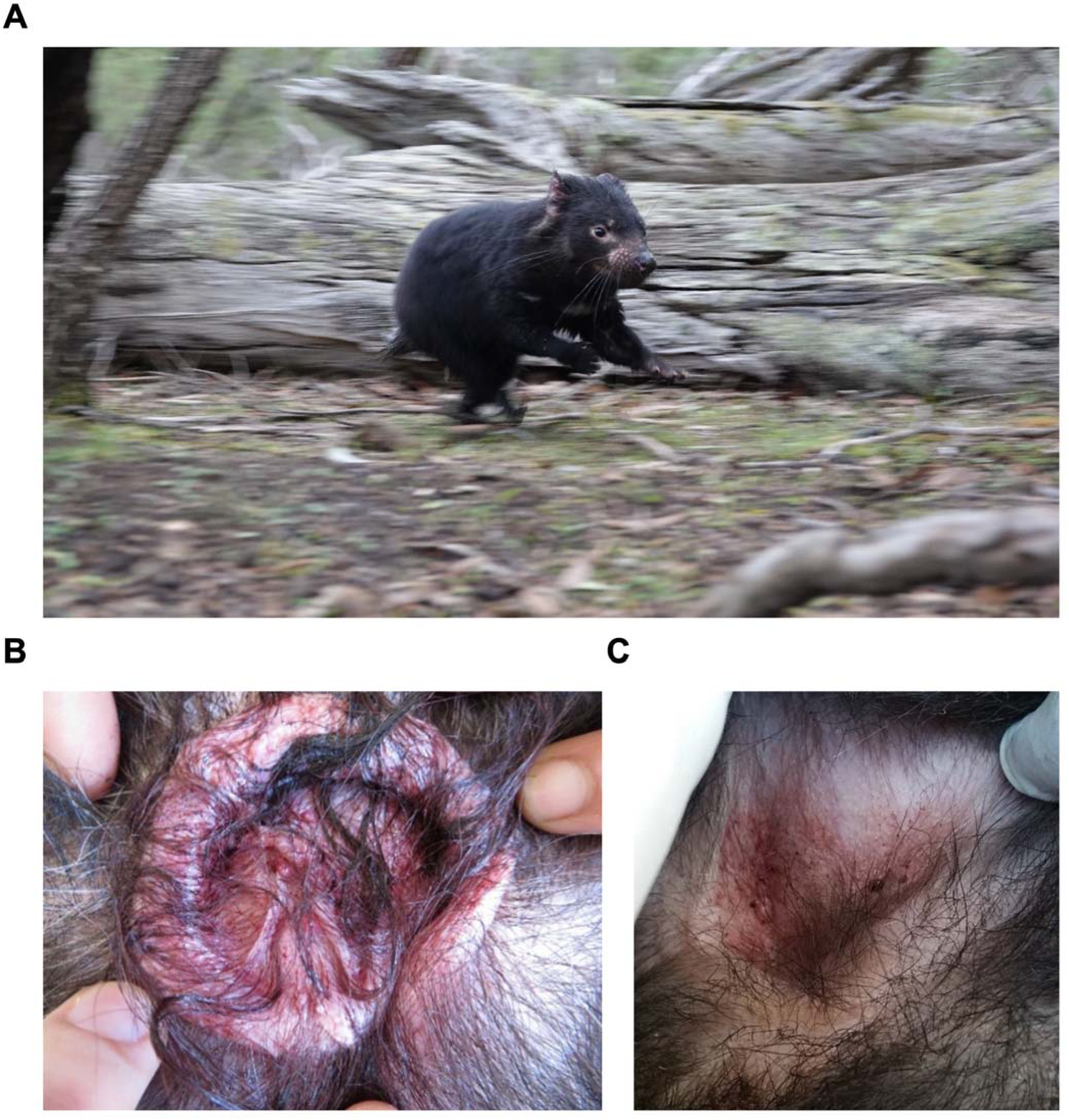
A) Image of a Tasmanian devil, B) Image of a pouch during breeding season with the red pouch oil secretions, C) Image of an inactive pouch. Photo credit: C. Hogg.

Previous studies have demonstrated that the devil pouch microbiome undergoes pronounced shifts in composition during reproduction and lactation, including a marked reduction in bacterial diversity and abundance leading up to birth (Peel *et al*. 2016, Ockert *et al*. 2024). These studies, and others into the tammar wallaby (*Notamacropus eugenii*) and the quokka (*Setonix brachyurus*) show there to be a reduction in the number of Gram-negative bacteria prior to the birth of young and during lactation (Old *et al*. 1998, Charlick *et al*. 1981, Peel *et al*. 2016, Ockert *et al*. 2024).

Pouch secretions are thought to play a critical role in protecting immunological naïve marsupial young during early development. Antimicrobial peptides (AMPs) such as cathelicidins are expressed in the pouch tissues where they are hypothesised to modulate the microbiome and provide direct protection against infection (Yadav *et al*. 1972, Ambatipudi *et al*. 2008, Peel *et al*. 2016). Compared to humans, marsupials, including Tasmanian devils, possess an unusually large and diverse repertoire of AMPs (Belov *et al*. 2007, Daly *et al*. 2008, Peel *et al*. 2016, 2024, 2025). AMPs are natural antibiotic molecules that form part of the innate immune system, with major families including cathelicidins and defensins. In devils, numerous cathelicidins are encoded in the genome, some of which are specifically expressed in the pouch skin with demonstrated broad-spectrum antimicrobial activity, including activity against bacterial strains that occur naturally in the pouch microbiome (Peel *et al*. 2016). In other marsupials such as tammar wallaby, cathelicidins are expressed in the skin of pouch young from as early as one day after birth (Daly *et al*. 2008).

Research on marsupial pouch secretions remains limited, with only three prior studies conducted, all focused on tammar wallabies or koalas (*Phascolarctos cinereus*) (Bobek & Deane 2002, Baudinette *et al*. 200, Ambatipudi *et al*. 2008). In koalas, several peptides were identified, some showing activity against Gram-negative bacteria; however, further characterisation was limited due to poor sequence homology with existing databases (Bobek & Deane 2002). Interestingly, fractionation of tammar pouch secretions revealed no direct antimicrobial activity. Only a single peptide, eugenin, was detected two weeks after birth, and it may play a role in promoting immune cell proliferation in the pouch skin (Baudinette *et al*. 2005). A subsequent investigation identified several proteins in the tammar wallaby pouch, such as beta-lactoglobulin (a major component of milk) and dermicidin (an antimicrobial peptide from sweat glands), some of which were active against Gram-negative bacteria (Ambatipudi *et al*. 2008). While previous work provides evidence for immune and bioactive proteins in pouch secretions, koalas and tammar wallabies do not exhibit the distinctive and copious red oil secretions that characterise the Tasmanian devil pouch (Hesterman *et al*. 2008).

Tasmanian devil milk contains a diverse suite of immune-related proteins, highlighting the importance of maternal secretions in protecting developing young. At mid-lactation, when the joey first exits the pouch, 6.6% of all genes expressed in the milk are associated with immunity (Hewavisenti *et al*. 2016). These include lysozyme C, one of the top ten most highly expressed milk genes, which is known to degrade bacterial cell walls; immunoglobulins that recognise and neutralise antigens; AMPs such as cathelicidins, defensins, and S100 family proteins that directly kill bacteria and modulate the immune system; and immune receptors such as major histocompatibility complex (MHC) molecules and cytokine signaling proteins (Hewavisenti *et al*. 2016). By analogy, devil pouch secretions may be immunologically complex, functioning as an additional line of protection for altricial young.

In this study, we employ a proteomics-based approach to characterise Tasmanian devil pouch secretions during reproduction for the first time.

## Materials and methods

### Sampling & extractions

Samples were collected from female Tasmanian devils (estimated age 14 months, N=4; estimated age 2.2 years, N=1) at Fentonbury and Bronte, Tasmania, Australia in May 2022 during the Save the Tasmanian Devil Program’s annual monitoring activities (University of Sydney Ethics ARA 2022/2243; Lazenby *et al*. 2018). All females had some red oil present (pouch score = 3, N=4) or max, puffy lipstick ring with oil drops (pouch score = 4, STDP, 2012). Samples were collected using a FLOQ swab (Copan Diagnostics, USA) that was inserted into the pouch and moved in a circular motion for five turns, before being flash frozen in a nunc tube using a dry shipper in the field. Samples were stored at -80^º^C until extraction.

Swab tips were immersed in 1 ml of UTC buffer (8M urea, 2M thiourea and 4% CHAPS in 0.1M Tris-HCl, pH 8) in a low protein binding Eppendorf tube and vortexed repeatedly until mixed. Eppendorf tubes containing swab tips underwent ten 30 s sonication cycles in ice-cold water with a Soniclean Ultrasonic Cleaner 25HD (650 W, 43 kHz, Soniclean, Australia). The swab tips were vortexed and centrifuged to extract the initial supernatant, which was carefully collected. Subsequently, the swab tips were inverted and reinserted into their respective Eppendorf tubes, followed by a second round of centrifugation to recover any residual liquid. The additional extract obtained was combined with the initial supernatant for each sample.

Each swab extract was processed using two methods to improve protein identification: direct protein analysis (method A) and acetone precipitation (method B). A 500 μL aliquot of each sample extract was allocated to method A, while another 500 μL aliquot was used for method B. For method B, five volumes (2.5 ml) of ice-cold acetone (-20 °C) were added to the extract, followed by centrifugation at 4 °C to obtain a protein pellet. The resulting pellet was washed with cold 80% acetone, and centrifuged again at 4 °C. The protein pellet was then resuspended in 50 μL of UTC buffer (8M urea, 2M thiourea and 4% CHAPS in 0.1M Tris-HCl, pH 8) and vortexed and sonicated in short bursts to ensure complete solubilisation. Protein concentration of samples from both method A and method B were estimated by using Bradford protein assay (Sigma, Australia) on a Varioskan Plate Reader (Thermo Scientific, Australia) following the supplier’s guidelines for sample dilutions and the BSA standard curve. Proteins were digested following the method described by (Bose *et al*. 2019). Briefly, a volume containing 100μg of sample was loaded into Amicon 10kDa MWCO filters (Millipore, Sigma, Australia) with four technical replicates were carried out for each sample. A solution of 8M urea in 0.1 M Tris-HCl (pH 8.5) was used to rinse the protein retained on the filter. Disulfide bonds were reduced by treatment with 10mM dithiothreitol (100 μL). Cysteines were alkylated by using 25 mM iodoacetamide (100 μL) in 8M urea, and the filters were incubated in the dark for 20 minutes. The buffer was then replaced with 50 mM ammonium bicarbonate (pH 8.0) before trypsin digestion. On filter protein digestion was carried out using trypsin (Promega, Australia) at a 1:50 enzyme to substrate ratio. The resulting peptides were dried completely using a Savant SpeedVac concentrator (Thermo Fisher Scientific, USA).

### Information-dependent acquisition (IDA) MS for discovery proteomics

The dehydrated peptides were redissolved in 1% formic acid (100 μL), and the technical repeats were pooled into a single sample and stored at 4 °C until analysis. An aliquot (2 μL) of each pooled sample was separated and analysed on an Ekspert nanoLC 415 (Eksigent, United States) coupled to a TripleTOF 6600 (SCIEX, United States) as previously described (Colgrave *et al*. 2017). Peptides were first desalted for five minutes on a ChromXP C18 trap column (3 μm, 120 Å, 10 × 0.3 mm) at a flow rate of 10 μL/min with 0.1% formic acid, and then separated on a ChromXP C18 analytical column (3 μm, 120 Å, 150 × 0.3 mm) at a flow rate of 5 μL/min. The mobile phases used were A) 5% DMSO, 0.1% formic acid, and 94.9% water; and B) 5% DMSO, 0.1% formic acid, 90% acetonitrile, and 4.9% water. A linear gradient of 3% to 25% solvent B was applied over 68 minutes, followed by a gradient from 25% to 35% of B over five minutes. This was followed by a rapid increase to 80% over two minutes, a two-minute hold at 90% B, a return to 3% B within one minute, and an eight-minute re-equilibration. The HPLC eluent was directly couple to the DuoSpray ion source of the TripleTOF 6600 mass spectrometer. The ionspray voltage was set at 5,500 V, with curtain gas at 138 kPa (20 psi), and ion source gases GS1 and GS2 at 103 kPa (15 psi) and 138 kPa (20 psi), respectively. The interface heater temperature was maintained at 150 °C. The eluent from the HPLC was directly coupled to the DuoSpray source of the TripleTOF 6600 MS. The ionspray voltage was set to 5,500 V; the curtain gas was set to 138 kPa (20 psi), and the ion source gas 1 and 2 (GS1 and GS2) were set to 103 and 138 kPa (15 and 20 psi). The heated interface was set to 150 °C. To enhance sensitivity for low signal-to-noise ratio (SNR) peptides, gas phase fractionation was employed. Information-dependent acquisition (IDA) was performed in top 30 mode using two replicate injections per sample, targeting precursor ion ranges of m/z 350-435 and m/z 430-1250, respectively. A third injection was analysed using DDA over the full mass range (m/z 350-1250).

### Protein identification

Discovery data files originating from the same individual, processed using protein extraction methods A and B and with or without gas phase fractionation, were collectively searched using ProteinPilot v5.0.3 (Shilov *et al*. 2007). Searchers were conducted against a FASTA database comprising NCBI RefSeq sequences (GCF_902635505.1; 45,895 protein sequences retrieved on 29/04/2025), in-house translated Tasmanian devil immune transcripts (497 sequences). This combined organism database was further merged with the publicly available common repository of adventitious proteins) to enable contaminant identification. The search effort was set to “Thorough ID”. Alkylation by iodoacetamide was selected, and Trypsin was chosen as the digestion enzyme. For ID emphasis, biological alterations were allowed, and the instrument type, TripleTOF 6600 (SCIEX), was specified. Proteins were identified at a 1% global false discover rate (FDR) (Tang *et al*. 2008). The SCIEX protein alignment template was applied to standardise protein identifications across samples.

### Peptide identification

Trypsin cleaved peptides were identified from each sample using two complementary approaches: database searching with ProteinPilot v5.0.3 (Shilov *et al*. 2007) and *de novo* sequencing with InstaNovo Plus (Eloff *et al*. 2025). For the database-driven approach, the background database and settings outlined in the preceding section were similarly implemented. The reported peptides from the database driven approach were filtered based on 1% global FDR. Results from the InstaNovo Plus algorithm were filtered based on a 99% probability threshold, peptide lengths of 6-30 amino acids, mass accuracy, and traceability to the background database described earlier.

### Data analysis and visualisation

The protein identification patterns across samples were visualised in an upset plot created using UpSetR package (Conway *et al*. 2017). Proteins present across a minimum of three samples were used as the basis for subsequent protein-level analyses.

For a preliminary peptide centric assessment of functional protein composition in swab samples, trypsin cleaved peptides, identified through two complementary strategies and consistently present in three or more samples were selected, and functionally annotated based on the Gene Ontology Biological Process (GO:BP) annotations of their parent protein using Unipept (Gurdeep Singh *et al*. 2019).

GO:BP overrepresentation analysis was then conducted on the subset of proteins selected based on their shared occurrence across samples. Overrepresented pathways were identified with g: Profiler, applying Fisher’s one-tailed test and a g:SCS *adj. p*-value threshold of 0.05 (Reimand *et al*. 2007). Overrepresentation analysis was performed using a custom background database constructed from RefSeq GO:BP annotations and translated Tasmanian devil immune transcript annotations. Functional protein annotations were obtained using PANNZER2 (Koskinen *et al*. 2015), and the database was assembled using a custom R script. To investigate the association of pouch proteins with cellular structures and complexes, Gene Ontology Cellular Component (GO:CC) overrepresentation analysis was performed using the same protein list and selection criteria as in the GO:BP analysis (Reimand *et al*. 2007), with a background database constructed from GO:CC annotations (Koskinen *et al*. 2015). The results from the GO:BP and GO:CC overrepresentation analyses were visualised using separate lollipop plots, created using ggplot2 R package (Wickham 2016).

The protein-protein interaction network (PPI) for the preselected set of proteins was constructed using STRING, with *Sarcophilus harrisii* selected as the organism (von Mering *et al*. 2003). The generated PPI network was further analysed in Cytoscape v3.10.3 (Shannon *et al*. 2003). Main clusters within the PPI network were identified by using the MCODE app (Bader and Hogue 2003). Cluster wise GO:BP overrepresentation analysis was conducted on each cluster to find overrepresented pathways (Reimand *et al*. 2007). Neighboring node proteins were annotated using GO:BP terms to elucidate their functional relationships with the proteins within each cluster (von Mering *et al*. 2003).

## Results

### Exploring the proteome profile of the pouch swab samples

Across all pouch swab samples, a total of 403 protein groups were identified. Individual samples yielded 236, 270, 259 and 297 protein groups in samples 1, 2, 3 and 4, respectively (**Fig. 2**). Of these, 153 protein groups were consistently detected in all four samples, while 200 were present in at least three samples.

**Figure 2.**
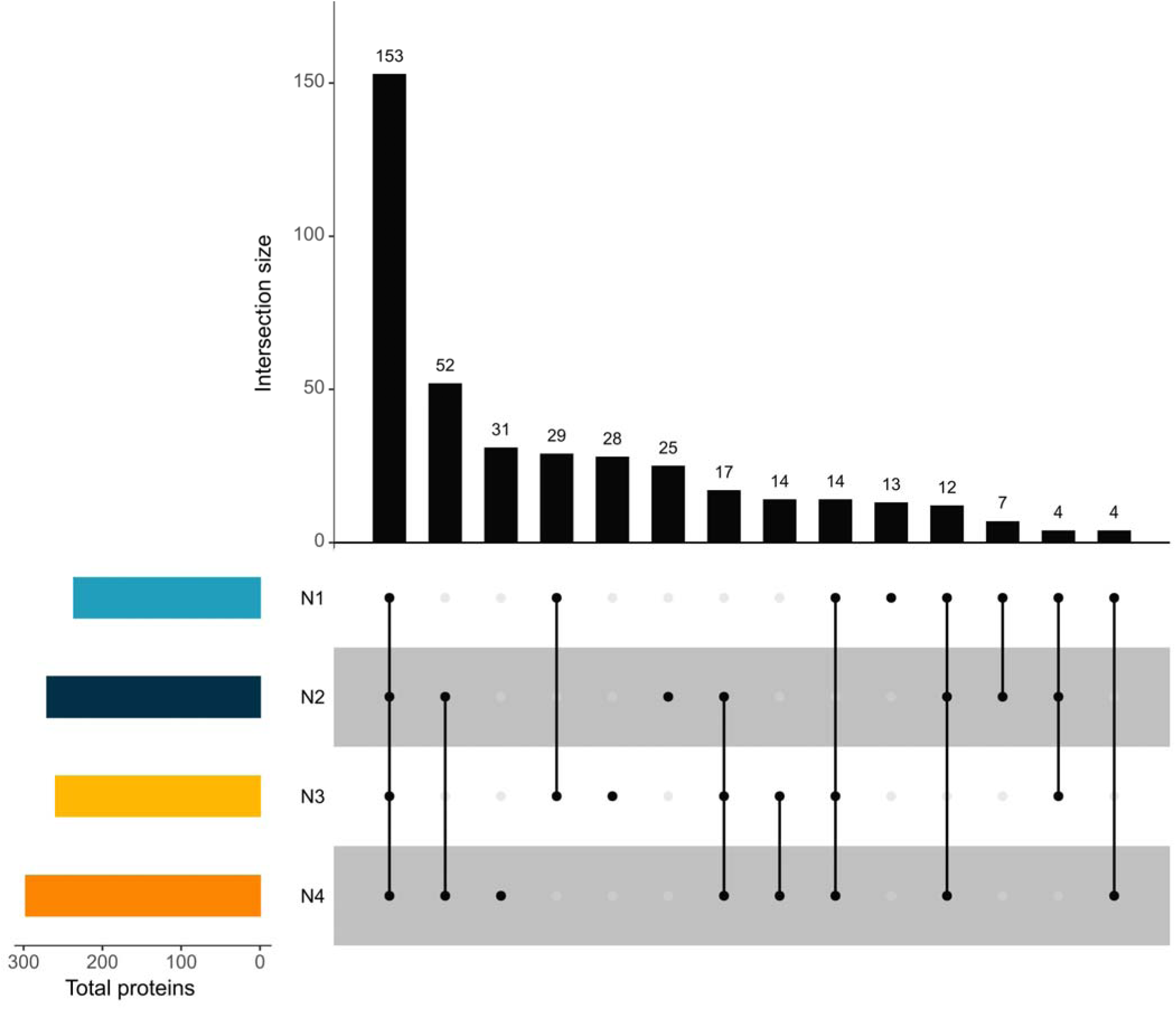
Comparison of the number of pouch proteins identified across four samples. The horizontal bar graph shows the total number of protein groups identified from each sample, while vertical bars indicate the number of overlapping proteins among samples. Intersecting black dots display the samples across which the proteins were shared (grey dot = not present).

### Peptide level evidence for proteins associated with key immune pathways

Gene Ontology Biological Process (GO:BP) analysis revealed multiple pathways with high peptide-level evidence, many of which are directly associated with immune function (**Fig. 3**). These immune pathways included iron-ion transport, defence response to Gram-negative bacterium, defence response to Gram-positive bacterium, killing cells of another organism, innate immune response, antigen processing and presentation of endogenous peptide antigen via MHC class 1b. Additional immune-related terms among the top-ranked pathways included adaptive immune response and antimicrobial humoral immune response mediated by antimicrobial peptides. Other highly represented biological processes included proteolysis, blood coagulation, oxidative stress response, keratinization, and vitamin D metabolism.

**Figure 3.**
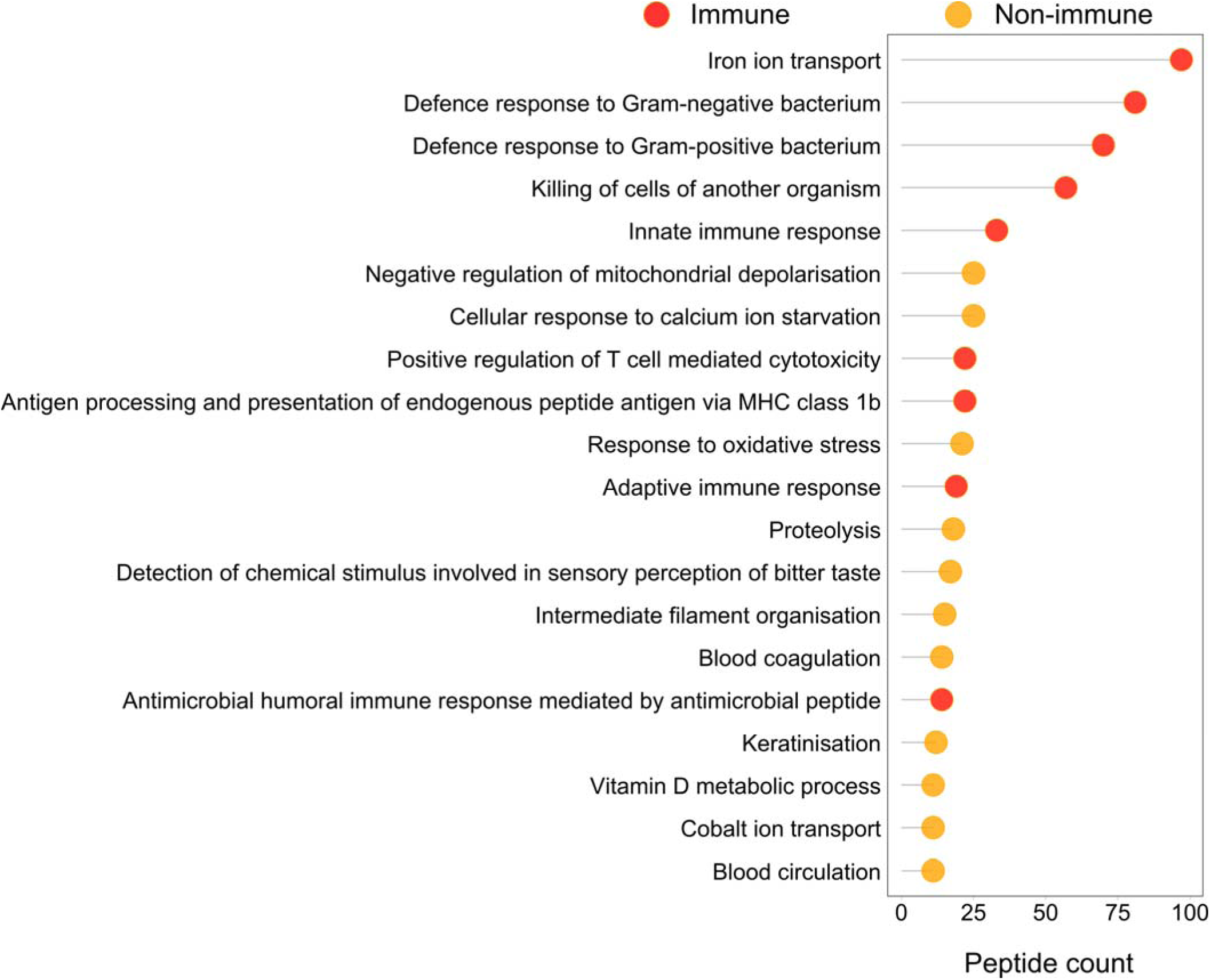
Most frequently detected GO:BP terms based on peptide evidence.

Extending the peptide-level observations, protein-level inference revealed a diverse array of immune proteins. Of the 200 proteins detected in at least three samples, 63 were associated with antimicrobial defence or immune regulation. These included immunoglobulins, components of the complement and coagulation cascade, antimicrobial proteins and peptides, inflammatory mediators, protease inhibitors, stress response proteins and S100 family proteins.

### Overrepresented immune pathways and protein localisation to immune complexes

GO:BP analysis of proteins shared across at least three samples indicated significant overrepresentation of immune pathways such as positive regulation of plasminogen activation, lymphocyte-mediated immunity, defence response to Gram-negative bacterium, antimicrobial humoral immune response mediated by antimicrobial peptide and adaptive immune response (**Fig. S1)**. Complementary GO:CC analysis showed strong overrepresentation of proteins localised to cellular immune complexes, including the pentameric IgM immunoglobulin complex, MHC class I peptide-loading complex, and lysosomes (**Fig. S2)**.

### Immune pathway overrepresentation persists within network clusters

Six primary clusters were identified within the PPI network, each showing distinct patterns of functional overrepresentation (**Fig. 4**). Cluster 1 showed overrepresentation of proteins involved in MHC class 1 peptide antigen assembly. Cluster 2 was overrepresented with acute-phase response proteins, while cluster 3 included proteins associated with lymphocyte mediated immunity and the adaptive immune response. Cluster 4 showed overrepresentation of proteins involved in the negative regulation of supramolecular fibre organisation and cluster 6 was overrepresented for proteins related to cellular oxidant detoxification. In contrast, cluster 5 did not exhibit any statistically significant functional overrepresentation.

**Figure 4.**
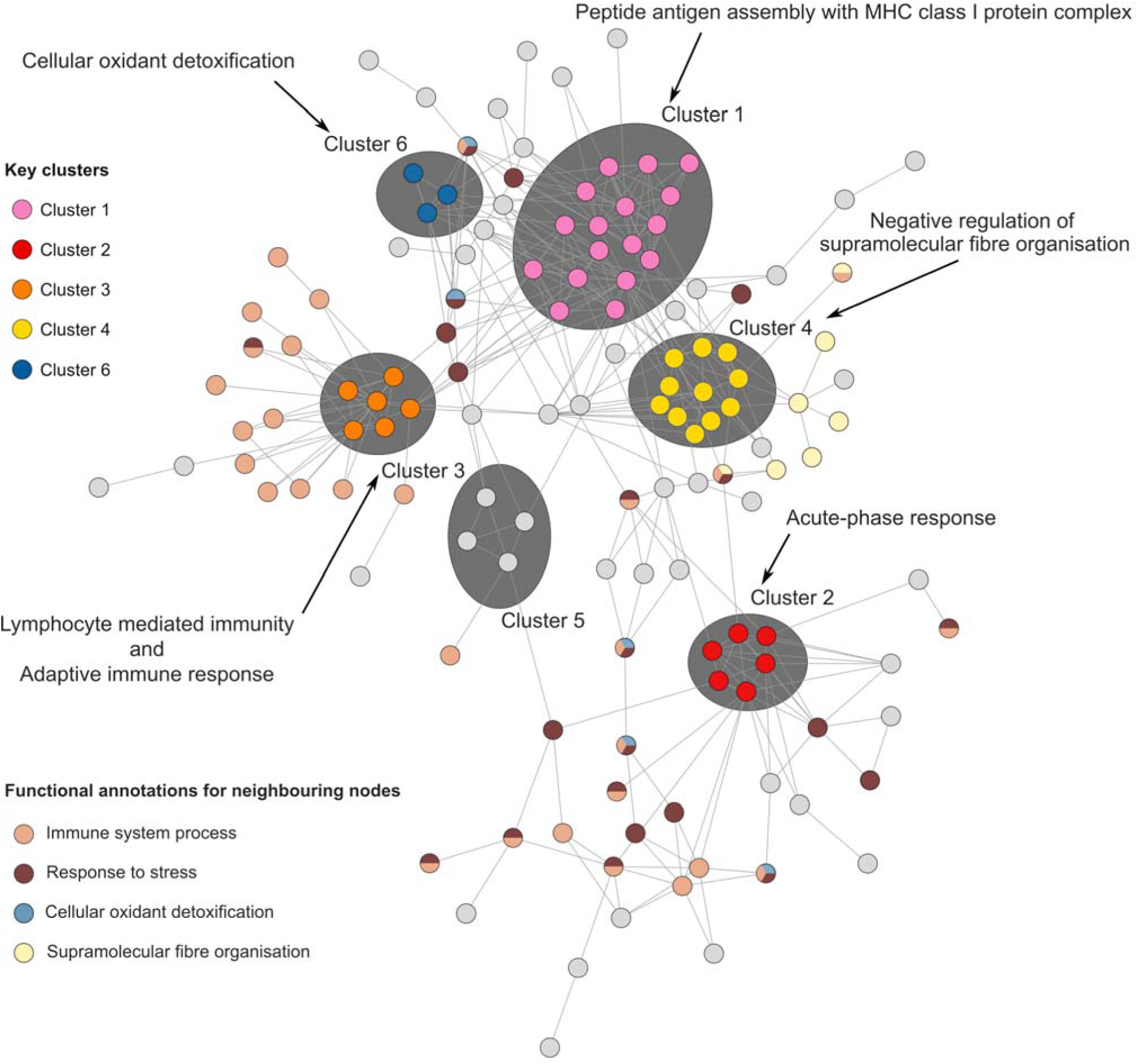
PPI network analysis of proteins shared across at least three samples. Clusters 1, 2, 3, 4 and 6 are colour-coded in pink, red, orange, gold, and blue, respectively. Neighbouring proteins with functional annotations related to immune system process, response to stress, cellular oxidant detoxification, and supramolecular organisation are colour-coded in light orange, brown, light blue and light yellow, respectively.

Adjacent nodes reinforced these functional associations, with proteins surrounding cluster 3 strongly linked to immune system processes and those neighbouring cluster 4 reflecting roles in supramolecular fibre organisation. Notably, all clusters were interconnected by stress response and immune system process proteins.

## Discussion

For the first time, we applied a proteomics approach to characterise the pouch secretions (“red oil”) produced by Tasmanian devils at the onset of the breeding season. This analysis addresses key knowledge gaps regarding pouch microbiome shifts between non-lactating and lactating females (Peel *et al*. 2016, Ockert *et al*. 2024). Our results reveal a complex protein profile consisting of proteins involved in the defence response to both Gram-negative and Gram-positive bacteria, in addition to proteins involved in both the innate and adaptive immune responses. These findings are consistent with previous observations of changes in the devil pouch during lactation and provides further details on pouch sections and protein pathways that are involved preparing the pouch for the altricial pouch young at birth.

Previous microbiome studies identified Actinobacteria, Bacteroidota, Firmicutes, Fusobacteria, and Proteobacteria as the dominant phyla, comprising 99% and 96% of the microbiome in lactating and non-lactating pouches, respectively (Ockert *et al*. 2024). Notably, the lactating pouch exhibited reduced microbial diversity and species richness, a pattern also observed in other marsupials such as koala (Maidment *et al*. 2023) and southern hairy nosed wombats (*Lasiorhinus latifrons*) (Weiss *et al*. 2021). Several genera known to contain pathogenic species, including *Vagococcus* (Pot *et al*. 1994) *Peptostreptococcus* (Higaki *et al*. 2000), *W5053* (Shi *et al*. 2022), *Ignatzschineria* (Deslandes *et al*. 2020)*, Moraxella* (Rabionet *et al*. 2016), *Wohlfahrtiimonas* (Schröttner *et al*. 2017), *Tissierella* (Caméléna *et al*. 2016), *and Clostridium sensu strico 1* (Cheung *et al*. 2023), were significantly reduced in lactating pouches. This microbial reduction was hypothesised to result from the presence of antimicrobial and immunomodulatory compounds secreted during lactation (Ockert *et al*. 2024). While previous studies indicated the possible presence of such compounds (Peel *et al*. 2016), the absence of proteomic level discovery left their functional expression unresolved.

The proteomic analysis of Tasmanian devil pouch secretions reveals a remarkable repertoire of immune and antimicrobial proteins, underpinning a complex local defence system akin to that described in other marsupials and mammals. Although previous transcriptomic studies have suggested the presence of such compounds, our proteomic data now provide direct evidence of their expression and functional relevance during breeding and lactation (Peel *et al*. 2016). Consistent with host defence requirements, the dataset is represented by proteins and peptides with activity against both Gram-negative and Gram-positive bacteria, including classic antimicrobials such as lysozyme (Cheng & Belov 2017), and multiple cathelicidins (Wang *et al*. 2011). Proteins in these pathways facilitate direct killing of microbes and can influence the composition of the pouch microbiota (Cheng & Belov 2017). Lysozyme, for instance, is a dominant antimicrobial protein found in animal secretions including marsupial pouch secretions that combines direct antimicrobial (Cheng & Belov 2017) and immunoregulatory functions—disrupting both bacteria and viruses and modulating the host’s immune response (Mann & Ndung’u 2020). Marsupial cathelicidins have been characterised as highly potent and resistant to salt, with essential roles in newborn immune protection and potential cross-protection against cancers such as devil facial tumour disease (Wang *et al*. 2011).

Among the most prominent findings are the presence of proteins involved in iron ion transport. Maintaining iron homeostasis is critical during infection, as both iron overload and deficiency can impair immune function and exacerbate pathogen load. Host proteins that sequester or transport iron, such as serotransferrin, limit bacterial access to this essential nutrient, thereby inhibiting pathogenic proliferation—a process termed “nutritional immunity” that is well-established in human and animal host defence (Cherayil 2011). We also identified key proteins implicated in the killing of foreign cells (e.g., complement proteins, lysozyme, cathelicidins) (Cheng & Belov 2017, Park *et al*. 2025), the innate immune response (e.g., acute-phase proteins, S100 family proteins) (Venteclef *et al*. 2011), and adaptive mechanisms such as antigen presentation via MHC class Ib complex (Bao *et al*. 2022) and T cell-mediated cytotoxicity (Ogg *et al*. 2019). Collectively, innate and adaptive immune pathway proteins ensure a rapid, non-specific response to invaders, followed by developed, antigen-specific responses, including immunological memory. Peptide and protein level evidence of T cell and B cell pathway components, including immunoglobulins IgA and IgM, highlight the spectrum of adaptive immunity available to developing joeys in an environment of high microbial challenge (Bonilla & Oettgen 2010).

Adaptive immunity is further supported by proteins integral to antigen processing and presentation, notably those involved in MHC class I and Ib pathways, critical for host responses against intracellular pathogens and for maintaining tissue homeostasis. This machinery is essential for the effective targeting of virally infected cells, other intracellular pathogen infected cells and tumours, again underscoring the pouch’s highly specialised immunological environment (Bonilla & Oettgen 2010; Ogg *et al*., 2019). Notably, cellular oxidant detoxification enzymes form part of a cohesive and supportive network. This network also includes proteins involved in the adaptive immune response such as MHC class I molecules and those mediating lymphocyte mediated immunity as well as components of the innate immune system, including acute-phase response proteins. These protein clusters are interconnected through stress response and immune system process proteins, reflecting critical inter-cluster cross talk that underpins both basic tissue homeostasis and rapid immune responsiveness. They help mitigate the potentially harmful effects of reactive oxygen species (Bassoy *et al*. 2021), maintain barrier function (Cheng & Belov 2017), and promote wound healing (Ramanathan *et al*. 2002), all of which are crucial in the context of the pouch’s dynamic, microbe-rich milieu (Cheng & Belov 2017). The prominent identification of a protein cluster involved in the organisation of supramolecular fibres suggests an additional dimension: the structural and renewal aspects of pouch skin (Ahn 2024) and how these may contribute directly to immune defence.

Due to their novel reproductive system, marsupials provide a unique opportunity to investigate the interplay between young development and peptides and proteins involved in their immunological protection. Here we have presented the first time the proteomic basis of Tasmanian devil pouch secretions, that when combined with previous transcriptomic work, start to provide further insights to how the altricial young survive and develop within the pouch environment. These findings position the marsupial pouch as a unique ecosystem, where robust immunological protection and tissue renewal converge to safeguard the highly vulnerable young during their earliest development.

## Author contribution statement

This project was conceived by CJH, with funding received to CJH, KB, MC. Sample collection was undertaken by CJH, protein analysis undertaken by MJM, AJ & MN-W, data interpretation was undertaken by MJM, EP, MC, CJH. The manuscript was written by MJM, EP, CJH. All authors edited and revised the manuscript.

## Acknowledgements

We acknowledge the traditional custodians, the Palawa people, of the lands upon which we collected the samples and pay respects to elders past and present. Thanks also to the field teams of the Save the Tasmanian Devil Program for their assistance in sample collection, namely Samantha Fox, Billie Lazenby and Drew Lee. Thanks also to Kimberley Batley and other members of the Australasian Wildlife Genomics Group.

