## Supplementary material for "Characterisation of pouch secretions from breeding Tasmanian devils"

**Corresponding author:**

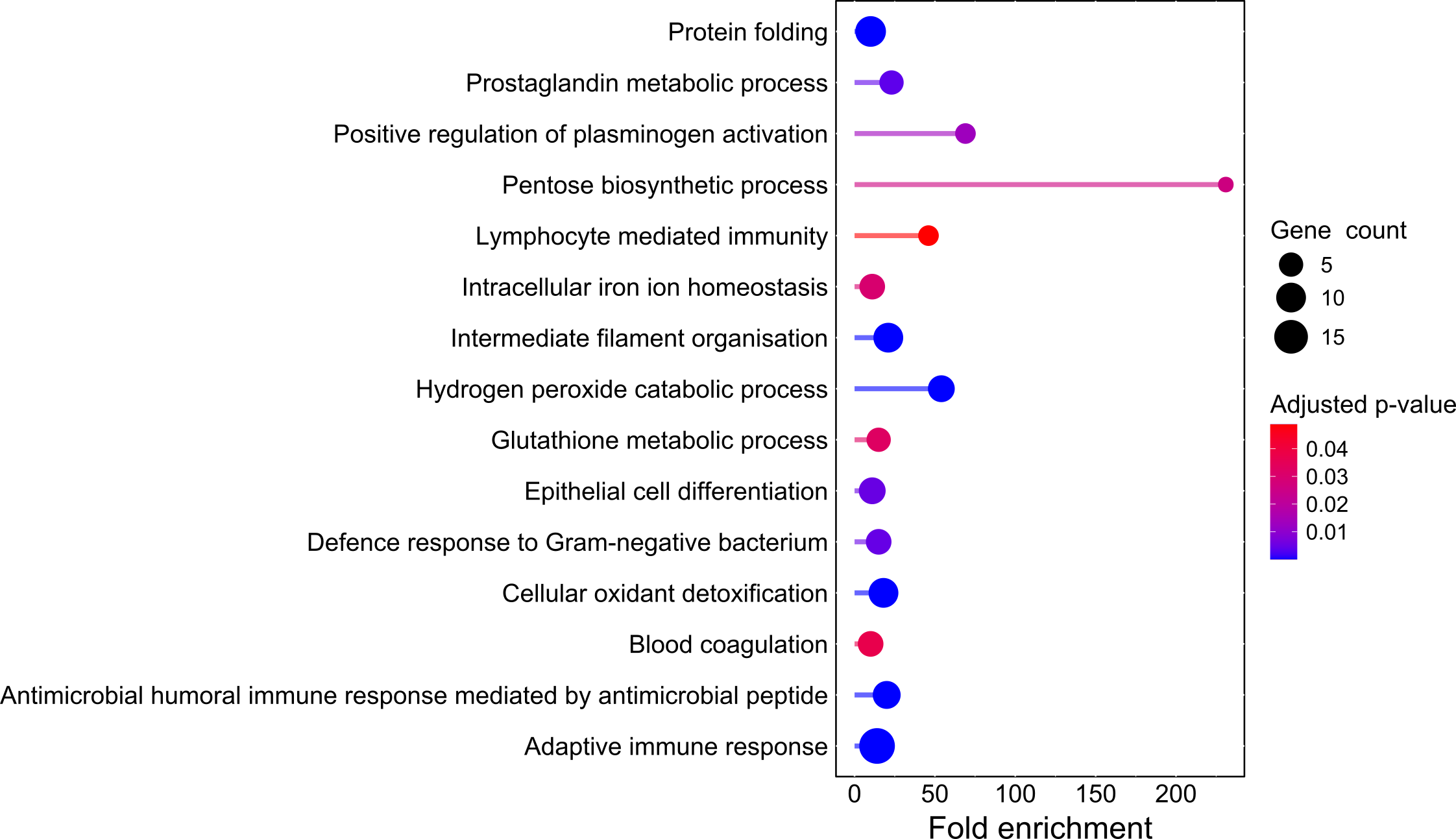

**Supplementary Figure 1** shows the overrepresented biological process terms across samples.

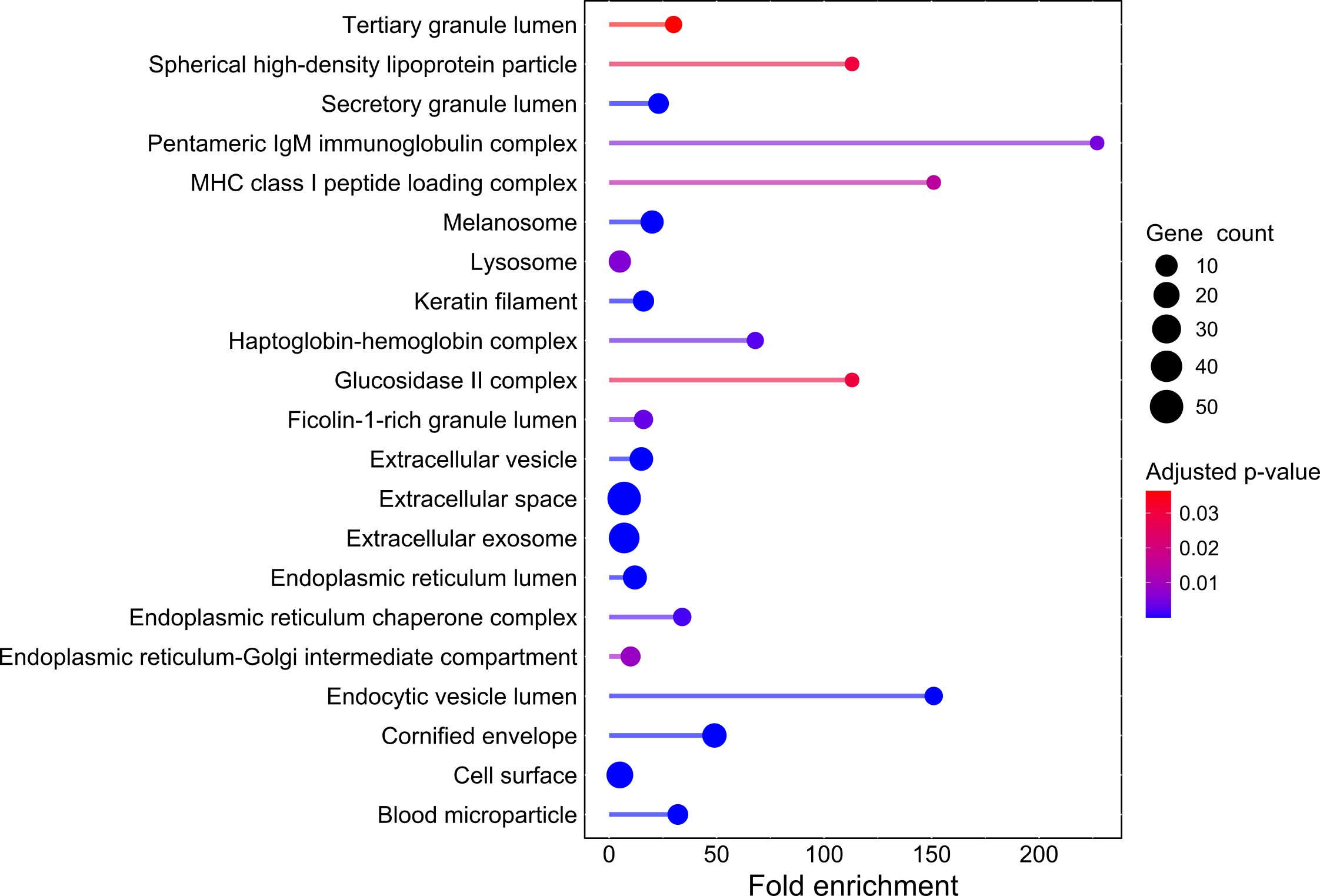

**Supplementary Figure 2** shows the overrepresented cellular component terms across samples.

**Supplementary Table 1** List of immune-related proteins identified in at least three Tasmanian devil pouch oil samples.

| **Representative accession** | **Protein name** |
| --- | --- |
| XP__031816989.1 | Ceruloplasmin-like [Sarcophilus harrisii] |
| XP__003768001.1 | Lactoperoxidase [Sarcophilus harrisii] |
| XP__003770901.1 | Lysozyme C [Sarcophilus harrisii] |
| XP__012403286.1 | Polymeric Immunoglobulin components receptor [Sarcophilus harrisii] |
| XP__031805230.1 | Pulmonary surfactant-associated protein D-like [Sarcophilus harrisii] |
| XP__031815616.1 | Serotransferrin [Sarcophilus harrisii] |
| XP__031807322.1 | Clusterin [Sarcophilus harrisii] |
| XP__031809570.1 | Double-headed protease inhibitor, submandibular gland-like [Sarcophilus harrisii] |
| EP_Saha_IGHA_IDMM | Immunoglobulin components |
| EP_Saha_IGHM_IDMM | Immunoglobulin components |
| XP__003768000.1 | Myeloperoxidase [Sarcophilus harrisii] |
| EP_Saha_IGKC | Immunoglobulin components |
| EP_Saha_IGHG_IDMM | Immunoglobulin components |
| EP_SahaCath5 | Cathelicidin-5 |
| XP__012398882.1 | Cathelicidin-3 [Sarcophilus harrisii] |
| XP__003762759.1 | Leukocyte elastase inhibitor [Sarcophilus harrisii] |
| XP__031816044.1 | Heat shock cognate 71 kDa protein [Sarcophilus harrisii] |
| XP__003758589.3 | Haptoglobin [Sarcophilus harrisii] |
| EP_Anst-IGHV3 | Immunoglobulin components |
| XP__023350610.2 | Cystatin [Sarcophilus harrisii] |
| XP__012401316.2 | Hemopexin, partial [Sarcophilus harrisii] |
| XP__003759347.3 | Glutathione peroxidase 3 [Sarcophilus harrisii] |
| XP__023354655.1 | Superoxide dismutase [Cu-Zn] [Sarcophilus harrisii] |
| XP__003768081.1 | Protein S100-A12 [Sarcophilus harrisii] |
| XP__031798513.1 | Peroxiredoxin-5, mitochondrial [Sarcophilus harrisii] |
| XP__031797811.1 | Cystatin-M [Sarcophilus harrisii] |
| XP__003772863.1 | Immunoglobulin components J chain [Sarcophilus harrisii] |
| XP__031825077.1 | Peroxiredoxin-1 [Sarcophilus harrisii] |
| XP__003773620.1 | Serum amyloid A protein [Sarcophilus harrisii] |
| XP__031805023.1 | Immunoglobulin components lambda-like polypeptide 5 [Sarcophilus harrisii] |
| EP_Anst-IGKV61 | Immunoglobulin components |
| XP__003768080.1 | Protein S100-A12 [Sarcophilus harrisii] |
| XP__003760881.2 | Complement factor D [Sarcophilus harrisii] |
| EP_Anst-IGKV94 | Immunoglobulin components |
| XP__023352649.1 | Myeloblastin [Sarcophilus harrisii] |
| EP_SahaCath6 | Cathelicidin-6 |
| XP__003769991.1 | Protein S100-A11 [Sarcophilus harrisii] |
| XP__012402110.1 | Beta-2-microglobulin [Sarcophilus harrisii] |
| EP_Saha_IGL_C2 | Immunoglobulin components |
| EP_Anst-IGKV4 | Immunoglobulin components |
| XP__031824437.1 | Complement component receptor 1-like protein [Sarcophilus harrisii] |
| EP_Anst-IGKV9 | Immunoglobulin components |
| XP__031804973.1 | Macrophage migration inhibitory factor [Sarcophilus harrisii] |
| XP__023355005.1 | Lymphocyte antigen 75 isoform X2 [Sarcophilus harrisii] |
| XP__031819170.1 | Kallikrein-7 [Sarcophilus harrisii] |
| XP__012409515.1 | Neutrophil gelatinase-associated lipocalin [Sarcophilus harrisii] |
| EP_Anst-IGHV5 | Immunoglobulin components |
| EP_Anst-IGKV64 | Immunoglobulin components |
| EP_Anst-IGKV19 | Immunoglobulin components |
| EP_Anst-IGKV38 | Immunoglobulin components |
| EP_Saha_IGL_C1 | Immunoglobulin components |
| EP_Anst-IGKV11 | Immunoglobulin components |
| EP_Anst-IGLV6 | Immunoglobulin components |
| EP_Anst-IGKV5 | Immunoglobulin components |
| XP__031803668.1 | Complement C3 [Sarcophilus harrisii] |
| XP__012406324.1 | Monocyte differentiation antigen CD14 [Sarcophilus harrisii] |
| XP__031816899.1 | Cathelicidin antimicrobial peptide isoform X2 [Sarcophilus harrisii] |
| EP_Anst-IGKV39 | Immunoglobulin components |
| XP__031809086.1 | Heat shock protein HSP 90-alpha [Sarcophilus harrisii] |
| XP__031819171.1 | Kallikrein-6 [Sarcophilus harrisii] |
| EP_Anst-IGKV111 | Immunoglobulin components |
| EP_Anst-IGKV13 | Immunoglobulin components |
| EP_Anst-IGLV1 | Immunoglobulin components |
